# From Quantitative Trait Loci towards Mechanisms: Linkage Integration Hypothesis Testing (LIgHT) Sheds Light on the Mechanisms of Genetically Modulated Stress Tolerance

**DOI:** 10.1101/2024.11.07.622539

**Authors:** Donghee Hoh, Isaac Osei-Bonsu, Atsuko Kanazawa, Nicholas Fisher, Jeffrey Cruz, Philip A. Roberts, Bao-Lam Huynh, David M. Kramer

## Abstract

The goal of this work is to assess the mechanistic bases of natural genetic variations in plant responses of photosynthesis to stress. To achieve this goal, we devised the Linkage Integration Hypothesis Testing (LIgHT) approach, comparing chromosomal locations of quantitative trait loci (QTL) for multiple phenotypes to distinguish between hypothetical mechanisms. As a use case, we explored genetic variations in photosynthesis-related processes under chilling stress in recombinant inbred lines of cowpea (*Vigna unguiculata* L. Walp.). We focused on photosynthesis-related parameters measurable in high throughput and indicative of proposed chilling responses, including the states of photosystems I (PSI) and II (PSII), photoprotective nonphotochemical quenching, PSII photodamage, and nyctinastic leaf movements (NLM). The patterns of QTL linkages indicated chilling stress tolerance is genetically controlled by avoiding PSII photodamage rather than PSI damage or NLM. This model was validated in a separate experiment measuring the rates of PSII photodamage and repair. Additional linkages suggest that chilling-induced damage to PSII is controlled by the thylakoid proton motive force and redox state of PSII. This regulation is modulated by thylakoid fatty acid composition, as suggested in Hoh et al., 2022. We propose the LIgHT approach can be broadly applied to test mechanisms underlying genetic variations.

**Highlight:** This work Introduces the Linkage Integration Hypothesis Testing (LIgHT) approach for mechanistic studies of natural variations, identifying photosynthetic regulatory mechanisms underlying natural variations to abiotic stresses by applying this approach.

## Introduction

Photosynthetic performance is strongly impacted by abiotic stress, accounting for substantial losses in plant productivity. Thus, improving this trait is critical for maintaining or expanding sustainable agriculture, particularly in a rapidly changing environment. Understanding how photosynthesis is regulated and contributes to yield under non-ideal conditions is critical to improving plant productivity. Stress resistance traits are thus the target of intensive efforts at breeding and engineering (Zelitch, 1982; Long *et al*., 2006; Raines, 2011). However, photosynthesis is a complex process sensitive to rapidly fluctuating combinations of temperature, water availability, light intensity, etc., that are not typically seen together under controlled laboratory conditions (Tikkanen *et al*., 2012; Cruz *et al*., 2016). Plants have adapted to meet the challenges of specific environments, and it may be possible to harness this natural genetic variation to improve crop performance in changing environments (Lawson *et al*., 2012; Theeuwen *et al*., 2022). However, such traits may not be present in our current crops or well-studied model systems in controlled environment laboratory settings. Thus, discovering the mechanistic bases of useful or adaptive photosynthetic traits will require exploring a wider range of genotypes and environmental conditions.

Recent studies showed strong linkages between single photosynthetic parameters in the field, especially linear electron flow (LEF) and the efficiency of photosystem II (PSII) photochemistry in light (Φ_II_), and seed yield under stress conditions such as drought in field experiments (Hamabwe *et al*., 2023; Keller *et al*., 2024; Taylor, 2024). Such results demonstrate that such field-based measurements can reveal associations between genomic variations and linkages between primary photosynthetic processes and yields.

Taking advantage of such natural genetic variations requires not only distinguishing between various limitations but also identifying which processes are controlled by genetic variations. This goal can be achieved by identifying statistical associations between measured traits with genetic polymorphisms in a panel or library of genetically diverse lines (Broman, 2001). Quantitative trait loci (QTL) and genome-wide association studies (GWAS) have been extensively used by plant breeders to identify genetic markers for desirable traits that can be used for introgression of these multiple traits into elite lines of crops (Boukar *et al*., 2016). For example, bulk or aggregated phenotypes based on the data, such as yield or disease resistance, were targeted for most QTL analyses (Muchero *et al*., 2013; Huynh *et al*., 2016). It has been much more challenging to identify the specific causative gene(s) associated with QTLs (Roff, 2007; Baxter, 2020), mainly because of the low genomic resolution of the methods (Miles and Wayne, 2008). Furthermore, it is difficult to determine the specific mechanisms of these associations based on any single photosynthetic parameter. For instance, decreases in Φ_II_ can result from low electron sink capacity, high levels of nonphotochemical quenching (NPQ), changes in leaf angle, or other physiological or morphological factors, each of which has multiple mechanistic causes.

Here, we provide an initial test of an approach we call “Linkage Integration Hypothesis Testing” (LIgHT), to distinguish such mechanistic connections by combining multiple photosynthetic and related parameters to provide “fingerprints” that distinguish among various mechanistic limitations. The increasing sophistication of high-throughput photosynthetic phenotyping, combined with powerful genetic approaches and biochemical methods, enables us to test for interactions among natural specific mechanisms that may underlie genetic variations in tolerance to low temperatures. Here, we combined data from high-throughput phenotyping tools that measure multiple mechanistically related photosynthetic phenotypes (Cruz *et al*., 2016; Kuhlgert *et al*., 2016; Theeuwen *et al*., 2022; von Bismarck *et al*., 2023).

As an initial use case, we focus on the extreme sensitivity of photosynthesis to even moderately low temperatures in *Vigna unguiculata.* L. Walp. (cowpea) (Hoh *et al*., 2022), a warm-climate species with a high level of genetic diversity in abiotic stress tolerance (Huynh *et al*., 2018). Chilling (or suboptimal) temperatures often significantly constrain photosynthesis, productivity, and geographical distribution of important cultivated crops (Allen and Ort, 2001). Counterintuitively, transient chilling (sub-optimal but non-freezing temperatures) can be a significant problem even with global climate change, which induces not only warming but also variations in temperatures, leading to unpredictable periods of increased and decreased growth temperatures (Gu *et al*., 2008). Multiple components of photosynthesis can be affected by chilling, including light reactions, carbon fixation, stomatal conductance and regulation of gene expression (Allen and Ort, 2001). Key steps in the light reactions have also been suggested to be the primary limitations under chilling, e.g. thylakoid electron transport, photodamage and repair of photosystem II (PSII) (Aro *et al*., 1993; Moon *et al*., 1995), photosystem I (PSI) (Sonoike, 1996; Shimakawa *et al*., 2024), activation of alternative electron sinks (Ivanov *et al*., 2012) and oxidative stress (Sassenrath *et al*., 1990; Hutchison *et al*., 2000). The primary limitations may be specific to different species, genotypes, developmental stages, or other environmental conditions.

Our recent work (Hoh *et al*., 2022) demonstrated a strong diversity of chilling tolerance in photosynthesis in cowpea recombinant inbred lines (RILs) that map to specific genetic loci and are linked to multiple photosynthetic processes and the contents of specific fatty acids.

Here, we further explore observed co-associations (or co-linkages) among multiple photosynthetic parameters rather than the parameters themselves to provide mechanistic fingerprints among photosynthetic parameters, allowing us to assess whether genetic variations in photosynthetic responses to stress are related to several distinct hypothetical mechanisms. The results suggest that a large fraction of the variations in photosynthetic efficiency were controlled by two QTL on chromosomes (Chrs) 4 and 9, associated with the control of the thylakoid proton motive force (*pmf*) and redox state of the primary quinone acceptor of PSII (Q_A_) and the PSII photodamage-repair cycle that determines the extent of photoinhibition. By contrast, we were able to eliminate an alternative hypothesis that the photosynthetic effects were caused by observed strong variations in nyctinastic leaf movements (NLM).

It is noteworthy that our hypothesis testing approach uses a statistical approach to test for co-associations among multiple parameters and does not rely on the identification of any specific polymorphism.

## Materials and Methods

### Plant materials

Cowpea RILs used for QTL mapping were selected by pre-screening nine pairs of RIL parental lines (Table S1). The selected population consisted of 90 RILs (F10 generation) originating from a cross between cultivar California Blackeye 27 (CB27) bred by the University of California, Riverside (Ehlers *et al*., 2000) and breeding line 24-125B-1 developed by Institute de Recherche Agricole pour le Développement (IRAD, Cameroon).

### Growth and Experimental Conditions

Cowpea seeds were planted in Suremix (Michigan Grower Products Inc, USA) with half-strength Hoagland’s nutrient solution and germinated under a 14hr: 10hr (day: night cycle) with a daylight intensity of 500 µmol photons m^-2^ s^-1^ and temperatures of 29 °C/19°C (day/night), 60% relative humidity in growth chambers (BioChambers, Winnipeg, Canada) (Figure S1A). Seedlings were then transferred to DEPI chambers and allowed to acclimate for one day under growth light and temperature conditions. Imaging of chlorophyll fluorescence parameters was initiated on the following days, with the light intensity changed every 30 minutes according to a 14/10 hour light/dark pattern based on a sinusoidal curve and a peak intensity of 500 µmol photons m^-2^ s^-1^ (Figure S1B), simulating a cloudless sunny day. On Day 2, day/night temperatures were shifted to 19 °C/13°C on the second imaging day for chilling treatment (Figure S1A). The temperatures were selected based on average field conditions from 2012 to 2016 in Tulare, Central Valley of California, where cowpea is usually grown in April, one month ahead of regular planting. Data is from the National Oceanic and Atmospheric Administration, https://www.noaa.gov (Table S2). Detailed experimental information for the temperature and relevant time points are referred to Table S3.

### Photosynthetic phenotyping

Chlorophyll fluorescence imaging was performed using Dynamic Environmental Phenotype Imager (DEPI) chambers (Cruz et al., 2016), with modifications described in (Tietz *et al*., 2017). Briefly, fluorescence images were captured in fully dark-adapted plants and at different times following illumination to obtain estimates of photosynthetic parameters using the methods described (Baker and Oxborough, 2004; Baker *et al*., 2007; Tietz *et al*., 2017). We measured values for steady-state (F_S_) and maximum fluorescence yields with Q_A_ fully reduced (F_M_’, F_M_”) collected after ∼0.3 s of saturating white light (∼10,000 µmol photons m^-2^ s^-1^) in light conditions or after 2 minutes of dark conditions, respectively. The oxidized Q_A_ state (F_0_’ and F_0_”) were collected immediately after 6 seconds of far-red illumination (approximately ∼4.6 µmol photons m^-2^ s^-1^) in light conditions or after 2 minutes of dark conditions, respectively. Those values were used to estimate Φ_II_, redox state of Q_A_, rapidly (q_E_) and slowly (q_I_ and q_Z_) relaxing contributions to NPQ. During the period of sinusoidal illumination, photosynthetic phenotyping was obtained two times per hour. Φ_II_ was derived from F_S_ and F_M_ images using previously reported methods (Cruz *et al*., 2016). Due to the large heliotropic movements of cowpea leaves, alternative equations (Tietz *et al*., 2017) were used for generating images of non-photochemical quenching (NPQt), photoinhibition-related quenching (qIt), energy-dependent quenching (qEt) and Q_A_ redox state PSII center opened (q_L_) rather than established methods for non-photochemical quenching use F_M_ and F_0_ images at the beginning of the day. All image processing was performed using Visual Phenomics 5 (https://caapp-msu.bitbucket.io/projects/visualphenomics5.0/), developed in-house using JAVA (Netbeans, www.java.com) and based on the open-source Fiji API (https://imagej.net/Fiji). Fluorescence and absorbance measurements were also performed using the hand-held MultispeQ V2.0, based on that described previously (Kuhlgert *et al*., 2016). To account for variations in leaf chlorophyll content, the light-induced thylakoid *pmf*, as estimated by the electrochromic shift (ECSt) parameter (Baker *et al*., 2007), was normalized to relative chlorophyll content.

### Linkage analysis and QTL mapping

Information on single nucleotide polymorphism (SNP) markers of genotype data of the CB27 x 24-125B-1 RIL population (Huynh et al., 2016) was followed by revised Chr numbering described by (Lonardi *et al*., 2019), based on expressed sequence tags (ESTs) produced by (Muchero *et al*., 2009; Lucas *et al*., 2011). IciMapping 4.1 (http://www.isbreeding.net) was used to construct the initial linkage map (Meng *et al*., 2015). Redundant markers were removed using the IciMapping “BIN’ function before constructing the linkage map. The linkage map was constructed using the Kosambi function using its RECORD ordering algorithm (Van Os *et al*., 2005), and then aligned against the cowpea consensus genetic map (Huynh *et al*., 2016). QTL analysis was performed using the Multiple QTL Mapping (MQM) model (genome scan with multiple QTL models), introduced by Ritsert Jansen initially (Jansen, 2004), as implemented in the Rqtl package (Broman and Sen, 2009). The *Rqtl fill.geno* function, based on a Hidden Markov Model, was used to fill in missing genotypic data. Levels of significance were determined using a permutation analysis implemented with the Rqtl *mqmpermutation* and *mqmscan* functions, over all replicates and with the number of permutations set at 1000 and a nominal significance cutoff of p < 0.05. The candidate genes in the QTL intervals were predicted by pseudomolecules (http://harvest-web.org) through BLAST in early-release genomes in Phytozome (www.phytozome.net/), and we developed Python codes for searching candidate genes. The candidate genes are annotated by Pfam, Panther, EuKaryotic Orthologous Groups (KOG), Kyoto Encyclopedia of Genes and Genomes (KO), Gene Ontology (GO), and best-hit Arabidopsis gene.

### Lincomycin treatment

For lincomycin experiments, detached leaves are vacuum infiltrated with 0.2 g/L lincomycin hydrochloride until complete inundation of cells by the solution. The control plants were vacuum infiltrated with deionized water (DI) using the same procedure. Infiltrated leaves were floated in plates with either lincomycin solution or DI water to avoid leaves’ dryness. Following infiltration, plates containing leaves and solution were kept under low light (50 µmol, m^-2^, s^-1^) for 20 min and then dark-adapted for 20 min to measure initial maximal PSII quantum efficiency (Fv/Fm). After that, Fv/Fm measurements were followed by 1hr of high light (HL) (1000 µmol, m^-2^, s^-1^) and 20 min dark adaptation to dissipate qE in the DEPI chamber. For the low-temperature treatment (LT), the temperature was decreased from 29°C to 19°C and 10°C every two hours of HL treatment (Figure S1C).

### Quantification of nyctinastic leaf movements (NLM)

Qualitative nyctinastic leaf movement (NLM) value measurements were obtained by measuring relative changes in the projected leaf tip-to-petiole distances of the time-resolved plant fluorescence images. Fluorescence images were taken during saturation pulses (i.e., used to estimate F_M_”), showing the strongest contrast against background interference. Each image was thresholded to separate the leaf area from the background using the triangle thresholding algorithm (Zack *et al*., 1977), which accounts for the vignetting effects of the cameras. The image regions for each plant were determined automatically by the code but verified manually, and the tip-to-petiole distance was taken as the long axis of a rectangle fitted to the projected leaf image. To account for differences in leaf morphology and size, fractional changes peak-to-peak distance normalizing to that of the presumed fully expanded and extended leaf states taken at midday.

## Results

### Effects of Chilling on Photosynthetic Responses and Leaf movement in Two Parental lines

To assess the mechanistic bases underlying natural genetic variations in photosynthetic responses to chilling stress, we initially screened nine parental pairs of cowpea RIL populations under chilling conditions (Table S1 and Fig. S1). Based on this screening, we selected the pair of CB27 and 24-125B-1, which showed the most pronounced differences in photosynthetic responses to chilling stress between the parental lines (Fig. S2).

Following the screening process, time series photosynthetic measurements were taken using dynamic environmental photosynthetic imaging (DEPI) (Fig. S3) and MultispeQ instruments (Fig. 1). The DEPI measurements were taken every 30 min for 14 hours for three days, while the MultispeQ measurements were taken five times per day to avoid disturbing the plants, as indicated in Fig. 1. MultisepQ measures not only chlorophyll fluorescence-derived parameters but also changes in absorbance by clamping the leaf. This capability enables observing more detailed photosynthetic responses (refer to Fig. S4 for hypothetical mechanistic contributors and measurable parameters of DEPI and MultispeQ). When the same measurements were made on both instruments, similar trends were observed. On Day 1 at control temperature (29°C/19°C) (CT), there were no or minimal differences between CB27 and 24-125B-1, but significant differences emerged under low temperature (LT) treatment (19°C/13°C) on Days 2 and 3 (Figs. 1 and S3).

**Figure 1.**
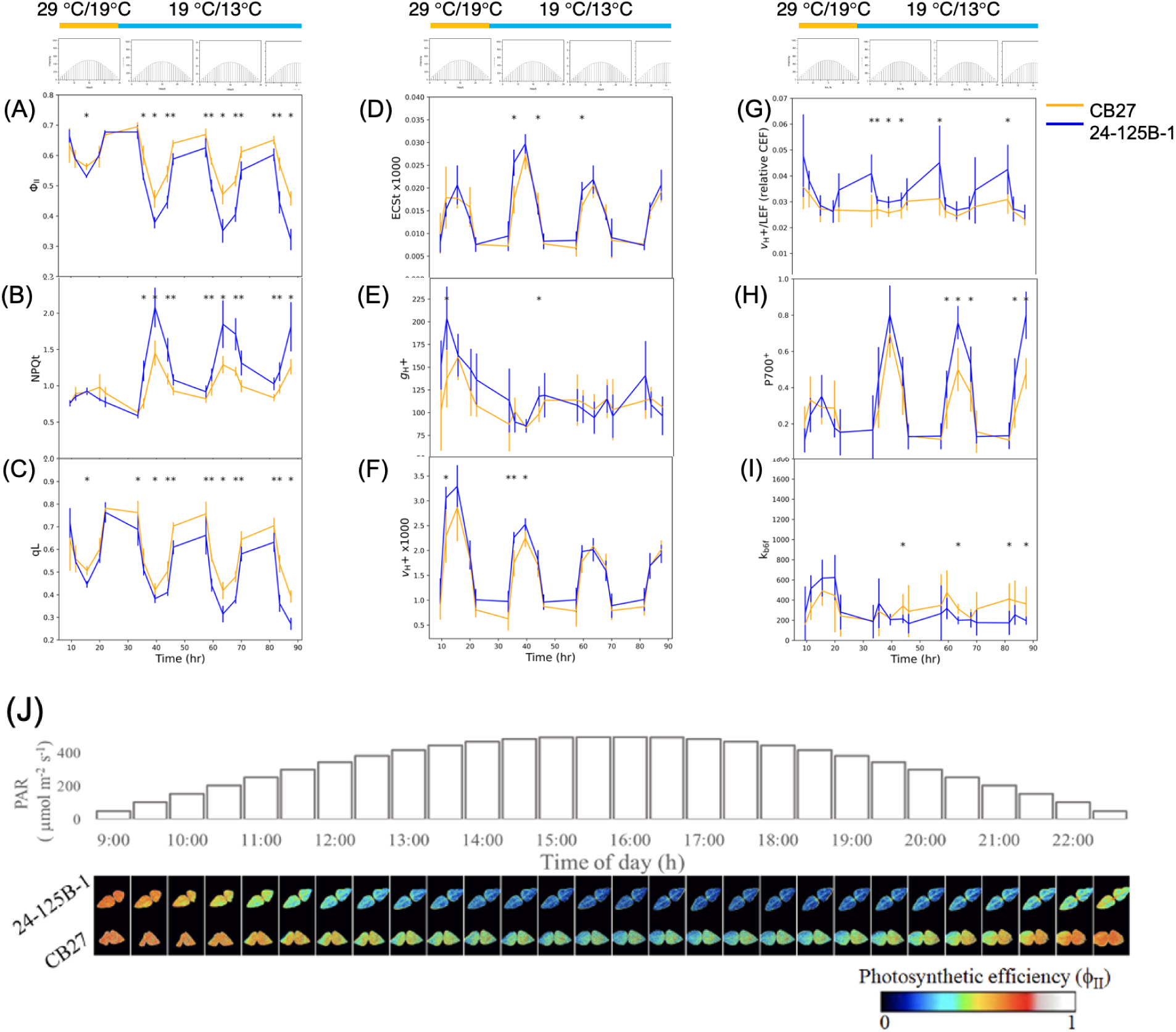
Effects of chilling on photosynthetic responses (A-I) and leaf movement (J) in two parental lines. (A-I) Time-resolved MultispeQ measurements of two parental lines at low temperature. (A, Φ_II_; B, NPQt; C, qL; D, ECSt (*pmf);* E; *g*H+, F; *v*H+, G: P_700_+; H; k_b6f_, I; *v*H+/LEF (relative CEF)). On Day 1, measurements were taken at the control temperature (CT, 29°C/19°C day/night temperatures, orange bars). In contrast, the following days were conducted at low temperature (LT, 19°C/13C°day/night temperature, blue bars). The patterns of light intensities (photosynthetically active radiation, PAR) and temperatures are displayed above each column of panels. The measurements were taken at five light intensities on Days 1-3, following a sinusoidal pattern: 103, 301, 500, 301 and 103 µmol photons m^-2^ s^-1^ (0.5, 2.5, 6.5, 11 and 13 hr after illumination). On Day 4, three measurements were taken at 103, 301, 500 µmol photons m^-2^ s^-1^ (0.5, 2.5, and 6.5 hr after illumination). The average response of n≥4 biological replicates for each photosynthetic phenotype value of two parental lines is orange for CB27 and blue for 24-125B-1. The significant differences between the two parental lines by t-test at each point are indicated as asterisks at the top of the plot (p<0.05). (J) Filmstrip view of sequential DEPI images that shows the changes in nyctinastic leaf movement (NLM) using false-coloring to reflect ɸ_II_ values throughout Day 2 of LT stress for the two parent lines. The light intensity in the DEPI chamber was increased by approximately 50 µmol photons m^-2^ s^-1^ every 30 min and images were captured at the same interval after each light intensity change over a 14-hour day. The top panel indicates the light intensity for each corresponding image.

In comparison to CB27, 24-125B-1 showed decreased Φ_II_ (Fig. 1A), increased NPQt (Fig. 1B) and decreased oxidation state of Q_A_ (qL) (Fig. 1C). These effects were accompanied by significantly higher ECSt, particularly at the beginning of days 2 and 3 (Fig. 1D), indicating a larger thylakoid *pmf*. However, the thylakoid proton conductivity, *g*H+, was either not significantly different or differed by only small amounts (Fig. 1E), implying that the increased *pmf* in 24-125B-1 could not be explained by slowing ATP synthase activity. The light-driven protons flux, estimated by the *v*H+ parameter, was increased in 24-125B-1, particularly at the beginning of Day 2, suggesting that the increased *pmf* was related to elevated proton fluxes (Fig. 1F).

The ratio of *v*H+/LEF can be used as an indicator of contributions to proton flux from cyclic electron flow (CEF) and LEF (Baker *et al*., 2007). In the absence of CEF, we expect a constant *v*H+/LEF because LEF should translocate a constant 3 H^+^/e^-^. Engagement of CEF should result in increased *v*H+/LEF. As shown in Fig. 1G, we observed periods of higher *v*H+/LEF in 24-125B-1, indicating that CEF likely contributed to the observed elevated *pmf* in 24-125B-1 throughout Day 2 and the beginning of Day 3 and Day 4.

We observed significantly increased levels of oxidized P_700_+ in 24-125B-1 on Day 3 (Fig. 1H), accompanied by decreased rate constant for P_700_+ re-reduction (k_b6f_, Fig. 1I), consistent with a larger photosynthetic control (PCON) likely imposed by the higher ΔpH component of the thylakoid *pmf*.

Interestingly, we observed significant variations in nyctinastic leaf movements (NLM) in parental lines during analyses of the DEPI data. The differences in NLM under LT are readily seen in the example images in Fig. 1J, in which parent line CB27 showed strong paraheliotropism (leaves pointing up) in the early morning but fully opened within 4 hours of light. By contrast, 24-125B-1 remained nearly fully open under all conditions.

In summary, the comparison between two parental lines showed that chilling sensitivity in 24-125B-1 is attributed to induced CEF and increased *pmf*, accompanied by decreased qL and Φ_II_, increased P_700_ oxidation due to decreased rate of electron transfer to PSI (increased PCON), leading to photoinhibition. The observed differences in photosynthetic responses and leaf movements in parental lines may be mechanistically and genetically linked. Thus, we explored natural genetic diversity in an RIL population derived from these two parental lines through the LIgHT approach to elucidate the mechanistic links between these effects and discern which processes are genetically modulated chilling sensitivity. The following section introduces the LIgHT approach to address these questions.

### Exploring Natural Variations in Chilling Responses in Photosynthesis: Linkage Integration Hypothesis Testing (LIgHT) approach

We devised the LIgHT approach to test for various hypothetical mechanisms exploring responses in natural genetic variations. LIgHT integrates QTL mapping with additional co-linkage tests and validation through subsequent experiments. The LIgHT can be a powerful tool for addressing complex interactions between genetic components and mechanisms by generating hypotheses based on experimental observations of multiple phenotypes in natural genetic variations. The LIgHT approach follows three distinct steps (Fig. 2):

1. Measure a range of phenotypes reflecting potentially mechanistically related processes using high-throughput tools.
2. Perform statistical analysis with phenotypic and genotypic data to map QTL and then compare linkages to generate an unbiased hypothesis. Co-linkage indicates that processes are either controlled by the same genetic loci or mechanistically related, indicating influence between them.
3. Test hypotheses through subsequent detailed experiments.

**Figure 2.**
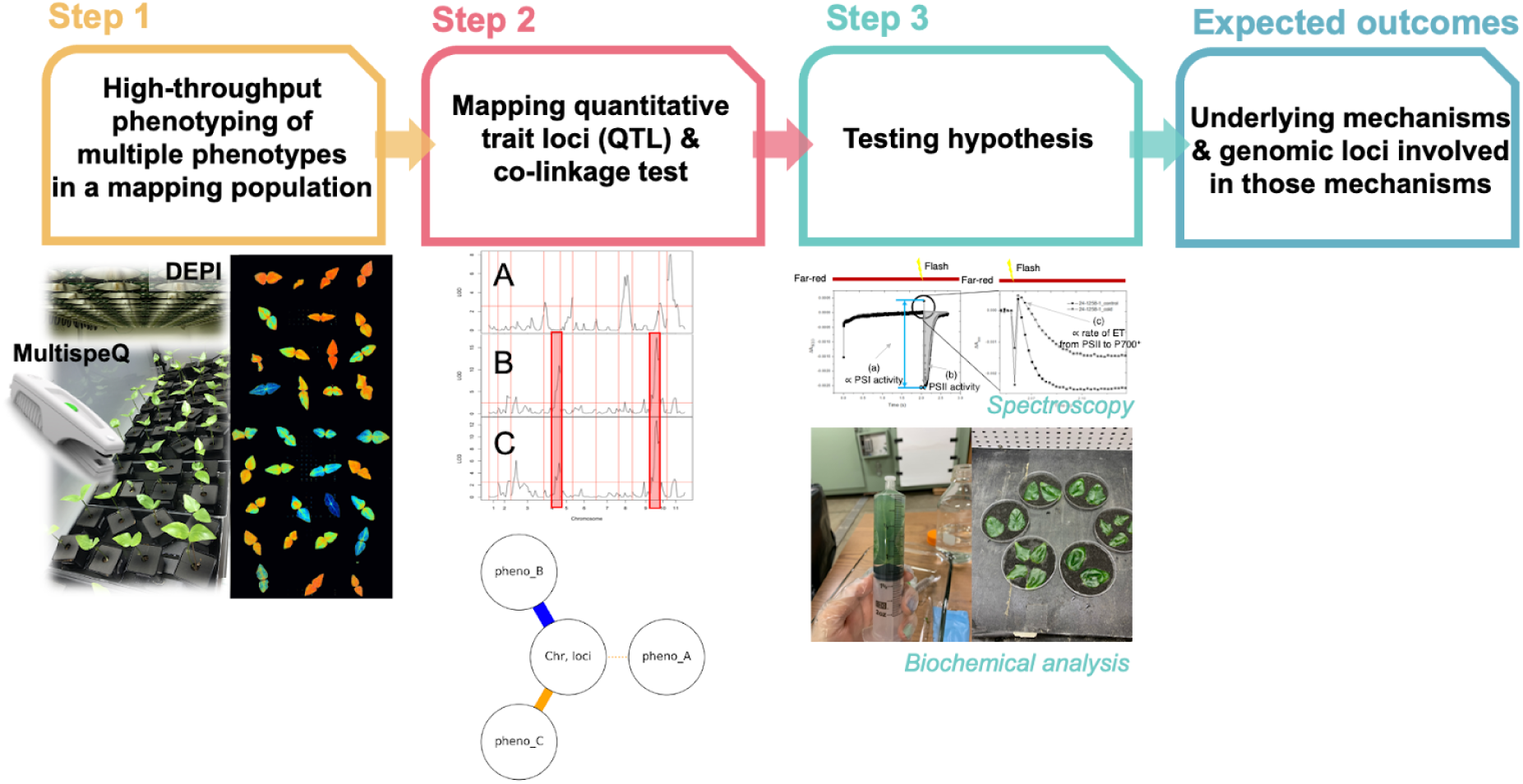
The schematic view of a general process of LIgHT. Step 1: Phenotyping. This step involves capturing multiple phenotypes that are mechanistically related by using multidimensional, high throughput, time-resolved, semi-simultaneous measurement tools in available germplasm, which has genetic diversity that can be mapped with genetic markers and phenotype data, such as RILs or diversity panels. **Step 2) QTL mapping and co-segregation test.** This step involves identifying QTL intervals and making comparisons. The presence of genetic or mechanistic linkage between distinct phenotypes is determined by comparing QTL intervals for multiple phenotypes. Overlapping QTL intervals in the same genetic locations indicate such a linkage (e.g., phenotypes B and C). “Linked” means either the same genetic loci control the processes or the processes are mechanistically related to the extent of influencing each other. If no linkage is observed (e.g., phenotypes A and phenotype B), genetic variation in phenotype A is not functionally or genetically linked to the variations in phenotype B. Based on the results of the co-linkage test, we can generate hypothetical models by comparing established mechanisms or directly generating hypothetical models. **Step 3) Testing Hypothesis.** This step involves testing the hypothesis generated from the QTL mapping and co-segregation tests through subsequent biochemical and biophysical experiments to identify specific and detailed underlying mechanisms.

We applied the LIgHT approach to explore potential primary chilling effects on photosynthetic responses, modulated by genetic diversity, in a subset of the data of the cowpea RIL population previously published (Hoh et al., 2022), with additional measurements. In this study, we extended the results and analysis from our previous work, incorporating additional measurements, genotypes, and analytical approaches to specifically distinguish among hypothetical mechanistic contributors to genetic variations in photosynthetic chilling responses. The following results sections will discuss each step of this approach and the results.

### Step 1 of LIgHT: High-Throughput Phenotyping of the RIL population

#### Dynamic Photosynthetic Responses and NLM in DEPI

Figure 3 shows the photosynthetic responses for the RIL population from DEPI, along with parent lines. It represents the time and genotype dependencies of photosynthetic parameters. Compared to CT, there were much larger genotype-dependent variations in measured parameters at LT (Figs. 3 and S5-6), with strong decreases in Φ_II_, qL, and qEt and increases in NPQt and qIt. Different from CT, which showed full recovery in Φ_II_ and NPQt by the end of the day, we observed a shift in the proportion of NPQ from qE (more rapidly recovering) to qI (more slowly reversible) components and strong decreases in qL at LT. These results imply that LT stress induces increased photoinhibition and a decrease in the rates of oxidation of Q_A_-that are not compensated by increases in NPQ, as discussed in (Kanazawa *et al*., 2021). We also observed a wide range of NLM phenotypes in the RIL population (Fig. S7). Some genotypes showed extents of motions that exceeded those seen in the two parental lines.

**Figure 3.**
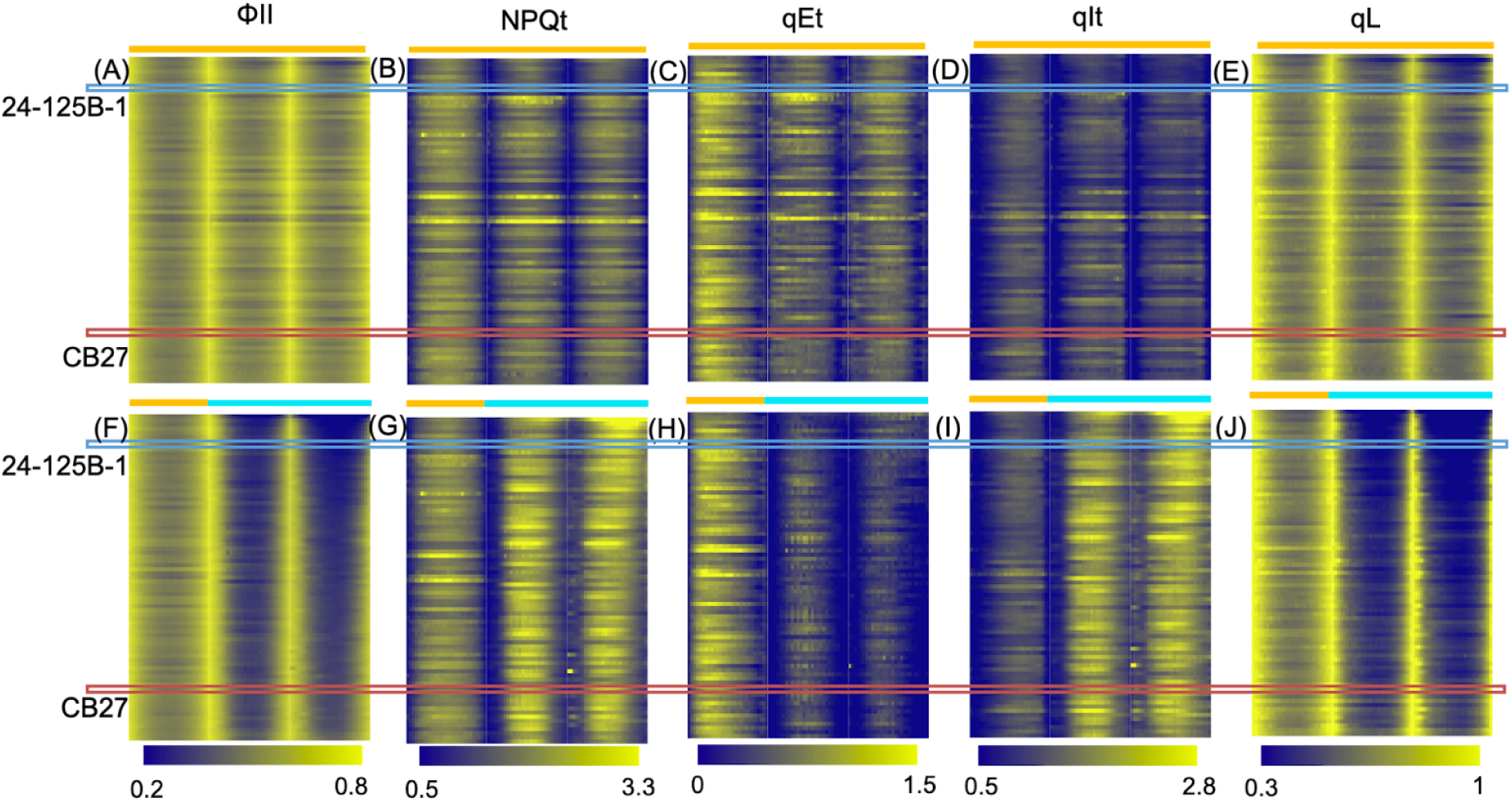
High-throughput photosynthetic phenotyping of recombinant lines (RILs) in DEPI chambers under control and low temperatures. Photosynthetic phenotyping of the CB27 x 24-125B-1 RIL population was conducted in a DEPI chamber on five-day-old seedlings over three days. Low (chilling) temperatures (LT) were applied under sinusoidal light on the second imaging day. Heat maps were generated using the OLIVER program (Tessmer et al. 2018) to display measured (non-normalized) averaged replicate values across the RIL population (n≥4). Each row denotes average values for a different genotype, with blue and red rectangles representing the two parental lines, 24-125B-1 and CB27, respectively. The remaining rows illustrate individual progenies within the RIL population. Five photosynthetic parameters were collected by the DEPI chamber every 30 minutes for 14 hours. The upper panels represent control conditions (A-E), while the lower panels (F-J) depict chilling conditions. The rows were ordered based on the average values of ΦII taken on Day 3 (the second day of chilling stress). Color legends for both conditions are standardized to allow for comparison. Significant changes in all parameters at low temperature compared to control conditions (day 1) are detailed in Figure S6.

#### Detailed Photosynthetic Responses using MultispeQ

We performed MultispeQ measurements across the entire RIL population to explore underlying mechanisms and potential genetic connections. The measurements were taken at a selected time in both temperature conditions in the middle of the third day of LT treatment (highest light intensity of the day). These measurements thereby capture the acclimatory responses to the different temperature conditions. Compared to CT, data from LT exposure show significant alteration in the mode of photosynthetic regulation processes. On average, compared to CT, an increase in CEF and a decrease in ATP synthase activity led to increased *pmf* and PCON, substantial increases in NPQ, and decreases in Φ_II_ and LEF at LT (Fig. S8). Strong variations were observed in these responses, likely reflecting genetic differences across the population.

### Step 2 of LIgHT: QTL Mapping and Co-linkage Test

#### Dynamic QTL intervals Associated with Photosynthetic Parameters and NLM

We identified distinct QTL intervals for each photosynthetic phenotype from DEPI and MulsitpeQ (Fig. S9-13). Interestingly, QTL intervals in Chrs 4 and 9 were observed for most photosynthetic parameters, as discussed below. However, the strongest QTL intervals for NLM were identified on Chrs 8, 10 and 11, specifically under LT conditions on Day 3 (Fig. S14). These intervals were strongest within about two hours after the start of morning illumination, when leaves were most rapidly transitioning from paraheliotropic to diaheliotropic positions.

#### Co-linkage testing Suggests PSII-related Limitations are the Primary Effect of Chilling Stress

We observed QTL intervals for most photosynthetic parameters in Chrs 4 and 9 and NLM in Chrs 8 and 10. Based on these findings, we test the co-association of DEPI, NLM and MultispeQ at a selected time point on Chrs 4, 9, 8, and 10. To effectively compare the co-linkages of multiple phenotypes within the identified loci, we developed co-linkage plots in the form of “Daisy Graphs.” These graphs feature specific loci represented by the center circles, encircled by varying phenotypes with the thickness of the connecting lines reflecting the logarithm of the odds (LOD) score for the association (Figs. 4 and S15). Solid lines within these graphs represent significant positive associations between the phenotype and the allele in the tolerant (CB27, pink) and sensitive (24-125B-1, light blue) lines.

**Figure 4.**
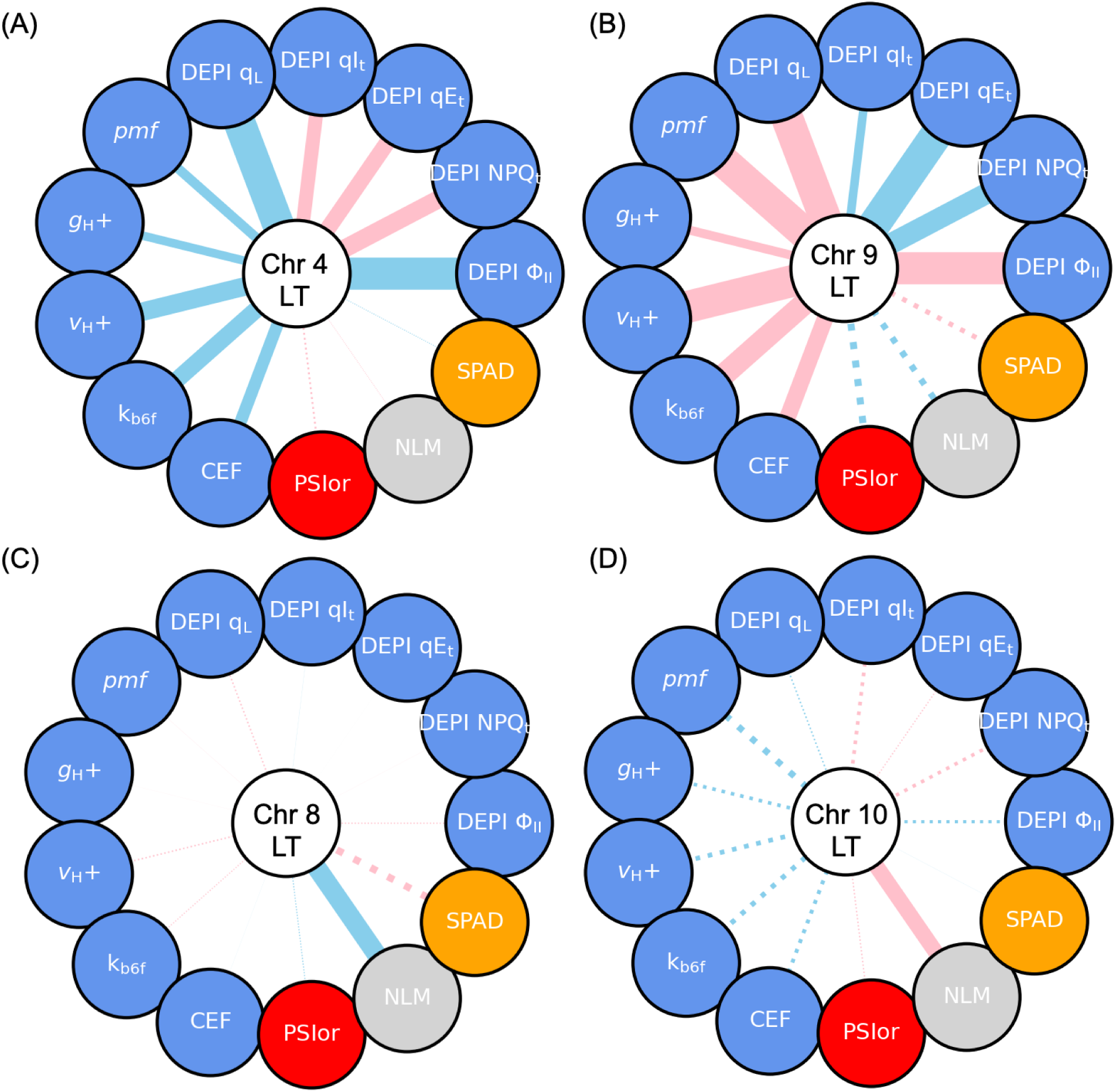
The Daisy plots show associations for selected QTL intervals of photosynthetic and non-photosynthetic parameters from DEPI and MultispeQ in LT at Chr 4, 59.04-64.45 cM (A) and Chr 9, 86.93-104.15 cM (B) Chr 8, 34.73-35.34 cM (C) Chr 10, 20-30.7cM (D). The LOD score plots from the previous figures (Figs. S9-14) have been reconceptualized in the form of “Daisy Graphs.” In these graphs, specific Chr is indicated in the center circles, and surrounding circles indicate different phenotypes. The thickness of the connecting lines is set proportional to the LOD score for association (Max LOD 10 is set to 10, so above the LOD 10 is shown as the same max thickness). The selected timepoint for photosynthetic parameters was at 1.5 hr before the end of Day 3 (206 µmol photons m-2 s-1). For NLM, 1 hour of illumination on Day 3 at 206 µmol photons m-2 s-1 was compared, as the effects were strongest within about 2 hours after the start of the morning illumination. Solid lines in the plots represent significant positive associations between the phenotype and the allele in the tolerant (CB27, pink) and sensitive (24-125B-1, light blue) lines. Phenotypes are color-coded by hypothetical mechanistic contributors to photosynthesis at low temperature: blue for PSII-related limitation, red for PSI acceptor side limitation, gray for NLM and orange for SPAD (relative chlorophyll content) (Fig. S4). Phenotypes with LOD scores below the threshold are shown as dashed lines. For more details on each plot, please refer to the original figures S9-14.

Within the contexts of observation in the parental lines, we tested three possible hypothetical mechanistic contributors: PSII-related limitation, PSI acceptor limitation and NLM (Figs. 1 and S4). In Chrs 4 and 9, we observed co-linkages for photosynthetic parameters related to PSII limitation but not PSI_or_ or NLM (Fig. 4 A and B). Interestingly, Chrs 4 and 9 have opposing effects on photosynthetic responses (See below). In Chrs 8 and 10, none of the photosynthetic parameters show linkages with NLM (Fig. 4 C and D). This result suggests that the effects of variations in NLM on photosynthetic properties were likely to be independent of those affected by QTLs on Chrs 4 and 9.

#### Time-resolved Co-association Distinguishes Causal links in Genetic Control of Photosynthetic Responses to Chilling

The co-linkage test of the LIgHT approach can facilitate the study of causal relationships when genomic loci overlap with multiple phenotypes across different time points. In principle, the appearance of QTL reflects the sequence of events leading to variations in phenotypes. Genetic variations causing earlier responses are mapped as QTL intervals showing large variations before those causing later responses. Thus, co-localized QTLs affecting traits at different time points suggest a causal link between the earlier and later-appearing traits.

In Chrs 4 and 9, on the first day of chilling stress (Day 2), the most pronounced response observed is appearance qEt intervals, indicating an initial response to chilling stress (Fig. 5 A and B). At LT, both Chrs showed a similar sequence where qEt, qIt, qL, and Φ_II_ appeared last, suggesting that qEt photoprotection occurs first and followed by photoinhibition and reduced Q_A_ (resulting in increased excitation pressure), leading to a decrease in Φ_II_ when stress exceeds the capacity of photoprotection. On Day 3, the responses differ slightly from those observed on Day 2, suggesting acclimatory or accumulatory responses to chilling stress on Day 3. Notable differences include ΦII appearing first and being sustained throughout the day and qL appearing much earlier than Day 2. The most significant difference between CT and LT in the Chr 4 and 9 intervals was the impact on NPQt and qIt (Figs 5 and S16). CT induced only a short, transient interval for qIt on Chr 4 in the morning on Day 3 (Fig. S16A) and none on Chr 9 (Fig. S16B).

**Figure 5.**
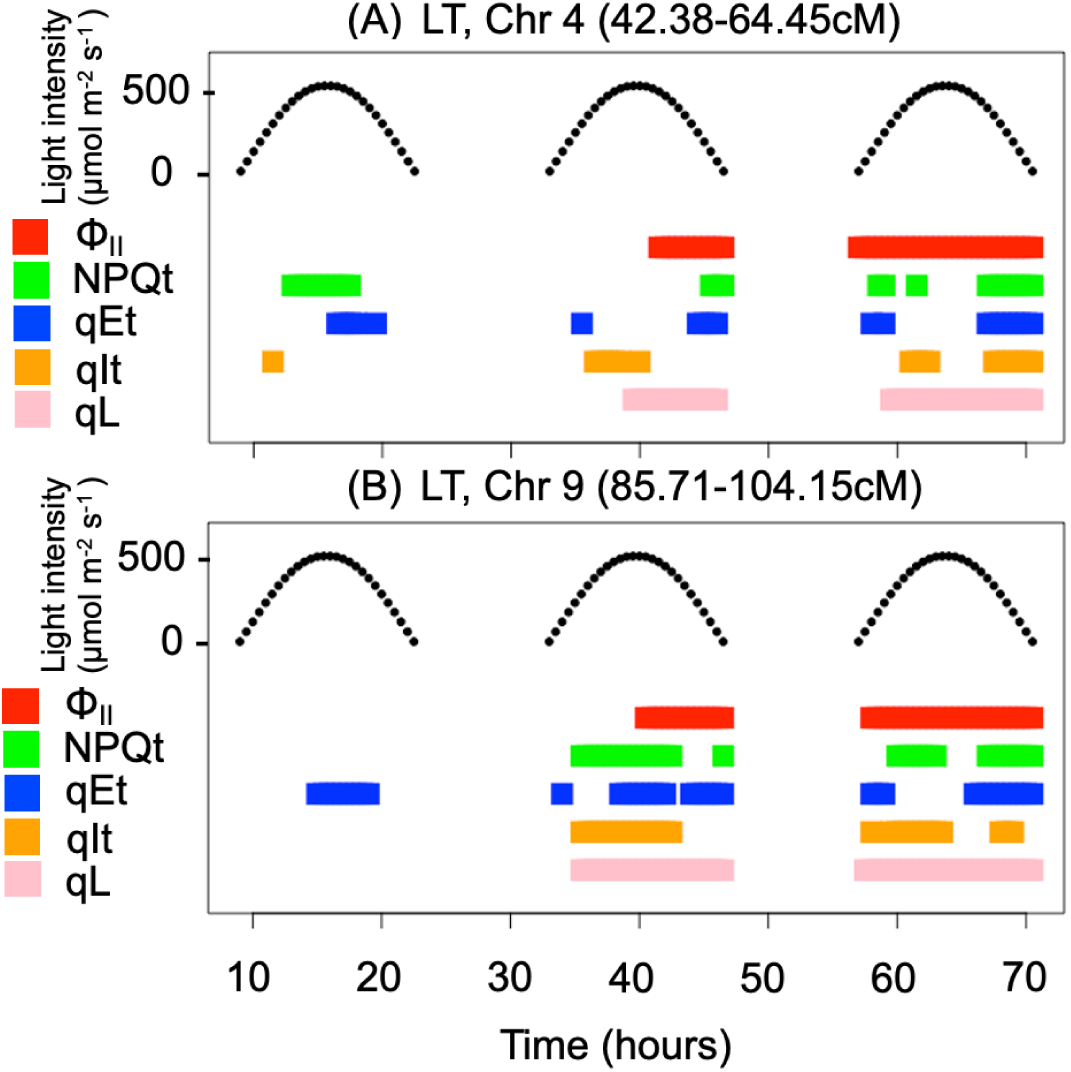
Time course for the appearance and disappearance of the QTLs for five photosynthetic parameters at selected genetic loci, Chrs 4 and 9 at LT. The appearance and disappearance of the QTL intervals for selected loci, Chr 4, 42.38-64.45 cM (A) and Chr 9, 85.71-104.15 cM (B) (CT is in Fig.S16). The time course for photosynthetically active radiation (PAR) is shown in the upper part of each panel. The appearance of significant QTL intervals for each phenotype is shown as filled rectangles with different colors: Φ_II_, red; NPQt, green; qEt, blue; qIt, orange; qL, pink.

In summary, the co-linkage test indicates that the primary effect of chilling is not due to PSI acceptor side limitation or NLM but rather due to PSII-related limitation. Therefore, we can rule out PSI limitation and NLM. Furthermore, time-resolved co-association patterns of QTL intervals were consistent with models where photoprotection occurred first and subsequently increased photoinhibition when stress exceeded photoprotection capacity. In the next section, we conducted tests focusing on PSII-related limitations to further validate the co-linkage test results.

### Step 3 of LIgHT: Genetic and Mechanistic Study of PSII Limitation Under Chilling Stress

So far, in Steps 1 and 2 of the LIgHT approach, we have narrowed down the primary effect of chilling stress to PSII-related limitations rather than PSI limitation or NLM. In this final step of LIgHT, we aim to assess genetically determined mechanisms underlying PSII limitation in more detail.

#### Genetic Effects on PSII Limitations in Response to Temperature Stress

In this section, we explore the genetic effects on the PSII limitation mechanism by analyzing the effect sizes and directionalities of genetic markers on the observed phenotypes, focusing on photoinhibition responses. Individuals of the RIL population are homozygous for each marker in the two parental lines, designated as either AA (allele from CB27, chilling tolerant) or BB (allele from 24-125B-1, chilling sensitive).

We first individually estimated genetic contributions from the QTL on Chrs 4 and 9 by dividing the population into two groups based on the presence of AA or BB markers for the QTL intervals. Consistent with what was observed in the Daisy plot, we found opposite effects of Chrs 4 and 9 (Fig. S17).

To test for additivity or epistasis, we assessed the combined effects of alleles by dividing the population into the four possible genetic combinations: AAAA, AABB, BBAA and BBBB. For instance, the AABB genotype has the CB27 allele on the Chr 4 QTL and the 24-125B-1 allele on the QTL for Chr 9. We observed polymorphisms within the QTL on Chrs 4 and 9 have additive but opposite effects on photosynthetic responses under both temperatures (Fig. S18-S19). These effects were more pronounced at LT, suggesting that the lower temperature accentuated the genotypic effects.

Next, we assessed genetic dependency on the relationship between two selected photosynthetic parameters to explore genetic effects on PSII limitation (Fig. 6). We found genetic effects on photosynthetic proton circuit and Q_A_ redox state modulate the responses to low-temperature stress (Fig. 6A), leading to PSII photoinhibition (Fig. 6B). Although average *g*H+ values across the population are lower at LT, they were similar across the genotypic groups at each temperature. Thus, the apparent lack of genetic contributions to *g*H+ appears to argue against a role in modulating ATP synthase activity at LT (Fig. S20).

**Figure 6.**
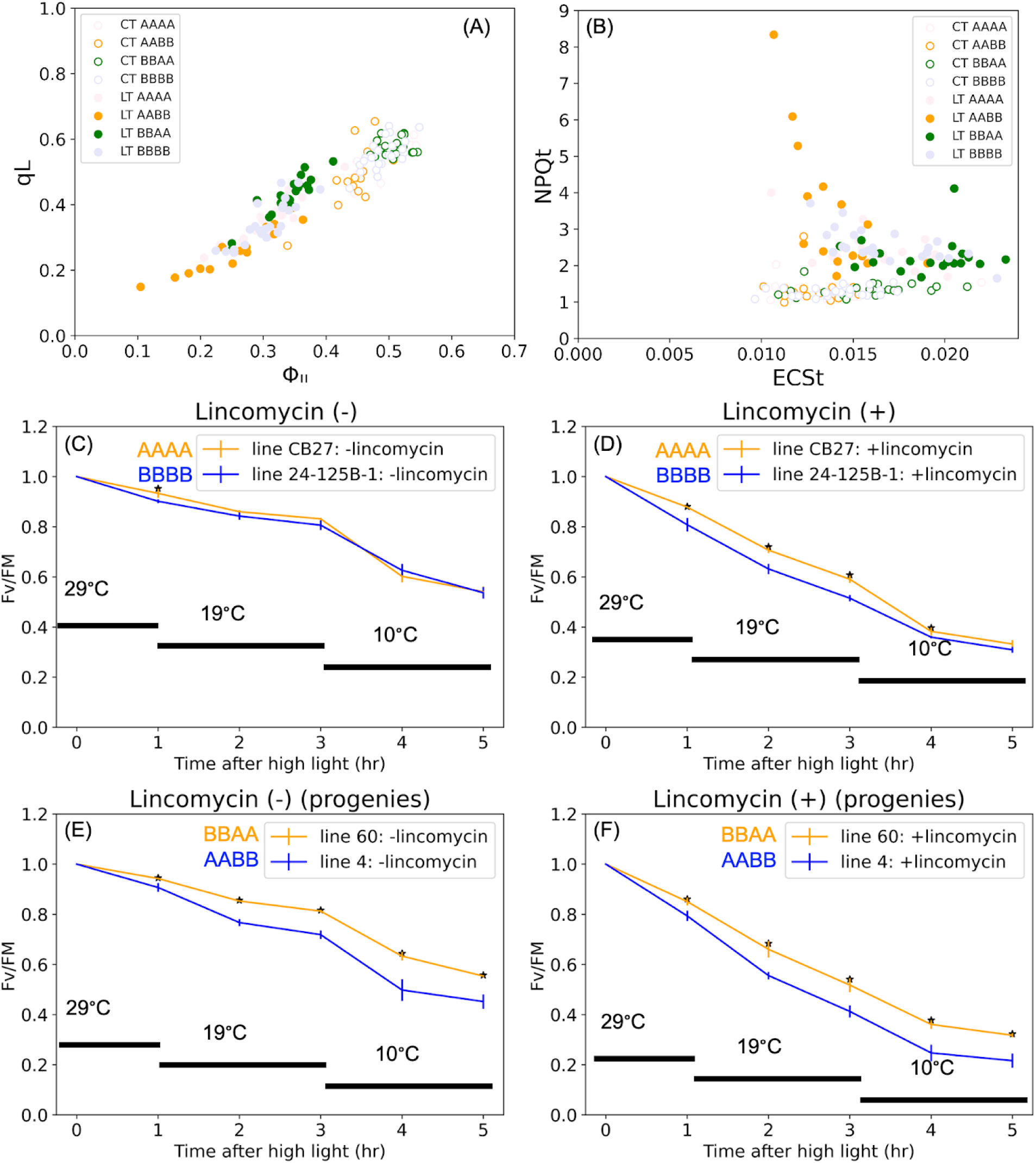
Genetically modulated PSII photoinhibition. **(A-B)** Relationships between photosynthetic responses grouped by different combinations of alleles for the identified QTLs in Chrs 4 and 9 under both conditions, CT and LT (CT: open circles, LT: closed circles). (A) **qL** versus **Φ_II_** from DEPI data, at midday on day 3 (highest light intensity, 500 µmol photons m^-2^ s^-1^). (B) **NPQt** versus **ECSt** from MultispeQ data, midday on day 4 (highest light intensity, 500 µmol photons m^-2^ s^-1^). AAAA and BBBB allele groups are indicated by light pink and light purple, respectively, while AABB and BBAA are colored orange and green, respectively. Detailed statistical analyses testing for differences in phenotypes between the allele groups are shown in Fig. S21. **(C-F)** PSII photodamage and repair during exposure to high light at a range of temperatures. Relative changes in the quantum efficiency of photosystem II (PSII) are estimated by the saturation flash-induced increases in chlorophyll fluorescence, as described in Materials and Methods. Panels A and B compare the two parental lines, while Panels C and D compare two selected progeny lines (RIL-60:BBAA and RIL-4: AABB) with combinations of alleles for the QTL intervals on Chrs 4 and 9 that show the largest differences on photosynthetic responses at LT. Intact, detached unifoliate leaves were vacuum infiltrated with either 0.2 g/L lincomycin (Panels B and D) to inhibit PSII repair or deionized water as a control (Panels A and C). The leaves were floated on these solutions during exposure to high light to prevent drying. Measurements were conducted using the DEPI chamber described in Figure S2, with the leaves exposed to constant, high light (1000 µmol, m^-2^, s^-1^) for one hour at a sequence of temperatures: control or growth temperature (CT, 29°C), low temperature (LT, 19°C, as used in the DEPI experiments) and a much lower temperature (10°C). Values of F_v_/F_M_’’ were measured periodically after a 20-minute dark period to allow for relaxation of qE, and normalized to the maximum PSII efficiency measured in dark-adapted samples (F_v_/F_M_). Averaged replicates (n≥3) ± S.D are shown.

In the following section, we will explore the underlying mechanisms of increased photoinhibition and its genetic effects by measuring the rate of photodamage and the repair in selected lines.

#### Genetic Effects on Photoinhibition at Low Temperature are Predominantly Controlled by Altering Rates of Photodamage

The results above suggest that the major QTL polymorphisms impact photosynthesis under both CT and LT but have cumulative, substantial secondary effects on PSII photoinhibition only at lower temperatures.

Two basic mechanisms are proposed to control PSII photoinhibition: altering the rates of PSII photodamage and PSII repair (Aro *et al*., 1993; Murata *et al*., 2007). To distinguish between these mechanisms, we measured the effects of high light illumination (1000 µmol m^-2^, s^-1^) on maximal PSII quantum efficiency (F_v_/F_m_) in the presence and absence of lincomycin, which blocks PSII repair by inhibiting D1 protein synthesis in the plastid (Tyystjärvi and Aro, 1996) (Fig. S1C). Since the effects of the alleles in the QTLs for Chrs 4 and 9 partly compensated for each other, we compared the two parental lines (CB27 and 24-125B-1, Fig. 6C and D) with two selected progeny lines from the groups that show the largest differences in photosynthetic responses, RIL-60 (BBAA) and RIL-4 (AABB) (Fig. 6E and F).

In the absence of lincomycin, the parental lines show only minor differences in PSII efficiency loss during high light exposure (Fig. 6C). However, with lincomycin infiltration, the sensitive line (24-125B-1) showed stronger losses of PSII efficiency, which proportionally increased at lower temperatures (Fig. 6D). These results suggest that PSII was photodamaged more rapidly in the sensitive line, but that repair mechanisms maintained similar steady-state levels of PSII activity in both lines. Notably, RIL-4 and RIL-60 showed progressively larger increases in photoinhibition, with stronger effects in the presence of lincomycin. This result indicates that a substantial fraction of the increased photoinhibition was caused by increased rates of photodamage, with less contributions from repair.

## Discussion

Using rapid, high-throughput phenotyping tools and the LIgHT approach on a cowpea RIL library, we investigate the mechanisms within genetic variation that modulate photosynthetic responses to chilling stress through QTL mapping for the first time. This work has also elucidated the mechanistic linkages and causal relationships contributing to the plant’s relative sensitivity to chilling stress.

### The LIgHT Approach Identified the Non-Primary Effects of Chilling on Photosynthesis

The analysis of the RIL library under CT and LT conditions revealed genetically controlled variations in many photosynthetic processes. Three notable exceptions were ATP synthase activity (*g*H+, Figs. 1E and S20), PSI overreduction (PSI_or_) and NLM (Fig. 4). We did observe a general reduction in *g*H+ going from CT to LT (Fig. S8H), as one would expect if the capacities for electron and proton flow and assimilation (Kanazawa and Kramer, 2002), sink strength (Takizawa *et al*., 2008) or onset of limitations at triose-phosphate utilization (Yang *et al*., 2016) were decreased at the lower temperature (Allen and Ort, 2001; ORT, 2002). However, the effect was not significantly different in the two parent lines (Fig. 1E), nor did we observe strong linkages to genetic markers (Fig. S13E), suggesting that modulation of ATP synthase activity did not contribute to the differences in chilling sensitivities within the RIL population under our conditions. It is possible, though, that a different population could exhibit such variations affecting chilling tolerance.

The lack of effects on Y_NA_ is interesting in light of the proposal that PSI photodamage, related to over-reduction, is a major factor in chilling-induced photodamage damage in some species, notably *Cucumis sativus* (Sonoike, 1996; Shimakawa *et al*., 2024), and in mutants that lack the ability to activate PCON (Tikkanen *et al*., 2012; Takagi *et al*., 2016; Kanazawa *et al*., 2017). Despite being quite chilling sensitive, we did not see any strong evidence for PSI over-reduction (PSI_or_) in cowpea (Figs. 4 and S13L). Instead, we observed strong PCON (likely due to decreased k_b6f_) (Figs. 4, S8E, and S13H), which resulted in net oxidation of P_700_ (P_700_+, Figs. 4, S8D, and S13H), preventing the accumulation of electrons on PSI electron acceptors.

We also observed strong induction of NLM, specifically under LT (Figs. 1J and S7). It has been proposed that these may protect against chilling damage to photosynthesis in some species (Huang *et al*., 2012, 2014). However, we did not observe obvious linkages to processes we measured on Chrs 4 and 9, including long-term changes in NPQt (Fig. 4), arguing against strong impact, at least under our conditions. The strongest QTL intervals for NLM were identified on Chrs 8, 10 and 11, specifically under LT conditions on Day 3 (Fig. S14), with the strongest responses occurring about two hours after the start of morning illumination. Additional leaf movement-related QTL intervals were seen (e.g., on Chrs 7 and 9 in the afternoon of Day 3) (Fig. S14B) but appeared to be associated with nutation motions, related to differences in growth of the stems and thus were not explored in detail. It is interesting to note, however, that these intervals did not overlap with those attributable to NLM, suggesting that different genetic components control these nutation motions.

### The LIgHT Approach Identified Genetically Determined Chilling Tolerance Network

The apparent co-linkages of photosynthetic parameters to QTLs on Chrs 4 and 9 and the order of their appearance suggest a model (Fig. 7) where the control of the light reactions by these loci is associated with increased thylakoid *pmf* (Figs. 1D and S8G), attributable to the activation of CEF (Figs. 1G and S8K), which results in more reduced Q_A_ (Figs.1, S5 and S6) and oxidized P_700_+ (Figs. S8). While these effects are seen under both experimental temperatures, they appear to have secondary effects at LT, resulting in strong differences in photoinhibition (Figs. 6, S5, S6), mainly caused by increased rates of photodamage (Fig. 6). This results in a strong shift in the sensitive lines, from qE to qI as the major form of NPQ (Figs. 3, S5-S6). This increased photodamage rate is associated with a net reduction of Q_A_ (Figs. 3, S5-S6), consistent with models where increased PSII excitation pressure (Huner *et al*., 1998) and elevated *pmf* (Figs. 1D and S8G), both of which will increase the rates of recombination reactions within PSII, resulting in the production of toxic singlet O_2_ (Ivanov *et al*., 2012; Telfer, 2014; Davis *et al*., 2016), and we thus propose this effect as the major contributor to the observed differences in chilling sensitivity of the light reactions. Such a mechanism is also consistent with the order of appearance of the linkages we observed in the time-resolved DEPI experiments, where qEt, qL and qIt preceded effects on Φ_II_ and NPQt (Figs. 5 and S11), providing insights into a causal relationship. Our recent work also demonstrated that specific fatty acid are mapped to Chrs 4 and 9, where we found QTLs in photosynthetic responses, suggesting that the specific fatty acid, phosphatidylglycerol 16:1^Δ3trans^, modulates this photosynthetic regulation model. This result suggests that this LigHT approach can be applied beyond photosynthetic phenotypes, highlighting its broad applicability (Hoh *et al*., 2022).

**Figure 7.**
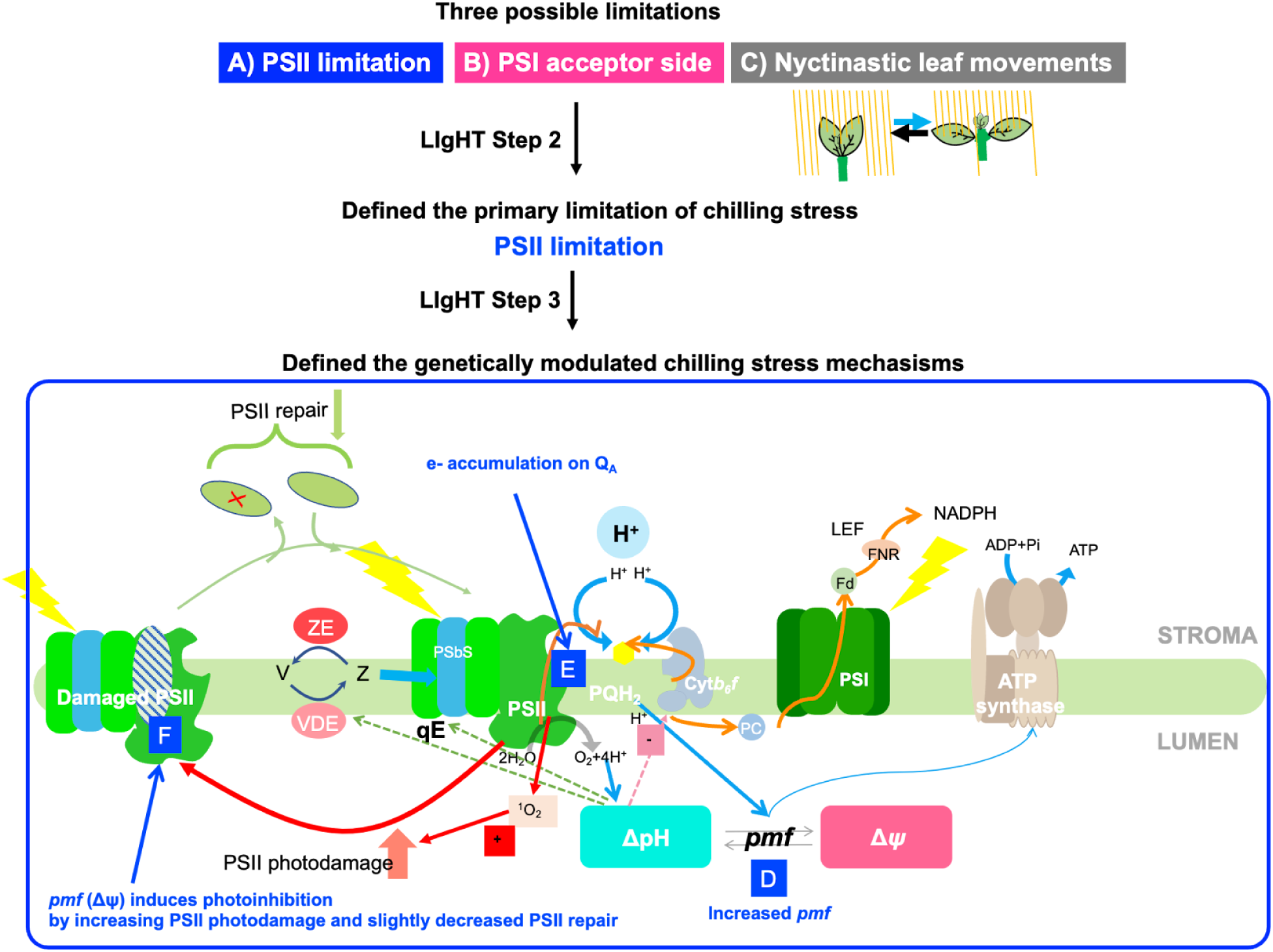
Summary of the LIgHT approach results and model, where genetically determined photosynthetic responses to chilling stress. Using high-throughput phenotyping and the LIgHT approach, we tested three main possible limitations: PSII limitation (A), PSI limitation (B), and NLM (C). Through the co-linkage test in LIgHT step 2, we confirmed that PSII limitation is the primary limitation under chilling stress, not PSI limitation or NLM. In LIgHT step 3, genetically modulated PSII inhibition mechanisms (Chrs 4 and 9) in response to chilling are defined. Chilling induces increased *pmf* (D), excitation pressure on Q_A_ (E), and photoinhibition (F) by substantially increasing photodamage with a minor decrease in PSII repair.

### Candidate Genes in QTL Intervals

We explored possible candidate genes in the minimum and maximum genetic range of QTL intervals in Chrs 4 and 9 that showed linkages for the photosynthetic parameters. First, based on the hypothesis that a common polymorphism will be responsible for the collective phenotypes associated with a QTL region, we determined which genes fell in the most likely intervals, encompassing the regions with LOD scores greater than the LOD significance threshold common to all the phenotypes. Second, because we cannot exclude the possibility that multiple polymorphisms contribute to the observed phenotypes, we explored the broader range of genomic regions that encompassed the full extent of all associations from any photosynthetic phenotype with LOD scores above the threshold.

For the QTL interval on Chr 4, we considered 13 overlapping QTL intervals ( 04-3-Φ_II_-LT, 04-3-NPQt-LT, 04-3-qEt-LT, 04-3-qIt-LT, 04-1-qL-LT from DEPI and 04-1-ECSt-LT, 04-1-gH+-LT, 04-1-LEF-LT, 04-1-P700+-LT, 04-1-Φ_II_-LT, 04-1-qL-LT, 04-1-vH+-LT, 04-1-kb6f-LT from MultispeQ), from which predicted the most likely common region as between 60 - 60.93 cM (flanking markers, 2_00148 and 2_07328), and contains 79 candidate genes (Table S6). One interesting candidate gene in this region is Deg1 (*Vigun04g188700*), a protease localized in the thylakoid lumen that is known to be involved in PSII repair by degrading damaged D1 (Kapri-Pardes *et al*., 2007) and OE33 subunits of PSII, as well as PC (Chassin *et al*., 2002). *deg1,* RNA interference transformed *Arabidopsis thaliana* with a reduced level of Deg1, is more sensitive to photoinhibition, showing accumulated D1 protein (inactive form) and less of its degradation products (Kapri-Pardes *et al*., 2007). A role for Deg1 in the modulation of the PSII repair cycle is consistent with our results, showing associations among Chr 4 genotypes, Φ_II_ and qIt (Fig.6) at LT, suggesting that certain alleles within this region show lower photoinhibition likely related to increase PSII repair. The broader (more inclusive) region of the Chr 4 QTL region, which extended between 34.47 - 64.45cM (flanking markers, 2_10801 and 2_04962) encompassed a total of 712 predicted coding regions (Table S7). This region included several additional photosynthesis-related genes, including the light-harvesting complex of photosystem II (LHCII) 5 (*Vigun04g167600*) and ferredoxin thioredoxin reductase (FTR) (*Vigun04g181000*), all of which could in principle contribute to chilling responses.

On Chr 9, we observed 14 overlapping QTL intervals (09-2-Φ_II_-LT, 09-2-NPQt-LT, 09-2-qEt-LT, 09-2-qIt-LT, 09-2-qL-LT from DEPI, 09-1-ECSt-LT, 09-1-gH+-LT, 09-1-kb6f-LT, 09-1-NPQt-LT, 09-1-P700+-LT, 09-1-qL-LT, 09-1-vH+-LT, 09-2-LEF-LT, 09-2-Φ_II_-LT from MultispeQ). The minimal common region was found to span 93.76-95.95cM (flanking markers, 2_11917 and 2_22085), containing 68 candidate genes (Table S8). One interesting candidate gene in this region is thioredoxin-h1 (trx-h1) (*Vigun09g249200*). Trx contains redox-active cysteine residues that are and reversely transfer the reducing potentials from light reactions to thiol-regulated enzymes. Trxs have conserved structures (WCGPC) to interact with target enzymes but react with different sets of target enzymes (Schürmann and Jacquot, 2000; Collin *et al*., 2003; Yoshida *et al*., 2014; Geigenberger *et al*., 2017). More than 20 isoforms of trx were found and categorized into several classes, trx f, h, m, x, y and z in the chloroplast (Collin *et al*., 2003; Yoshida *et al*., 2014; Geigenberger *et al*., 2017). Trx-h is eukaryotic trx and its potential target proteins are Triosephosphate isomerase, ADP-glucose pyrophosphorylase (AGPase) (Marx *et al*., 2003), peroxiredoxins (Rouhier *et al*., 2001; Marx *et al*., 2003; Maeda *et al*., 2004) and non-specific lipid transfer protein (Maeda *et al*., 2004). Ortiz et al. 2017 also identified trx gene (*Sb03g004670*) from a genome-wide association study (GWAS) of Sorghum with chlorophyll fluorescence and carbon assimilation measurements at LT (Ortiz *et al*., 2017). This region also contains genes for the large subunit of AGPase (APL2, *Vigun09g247600*), which is involved in starch biosynthesis and regulated by trx-h. A homologous gene in *Arabidopsis* (At1g27680) was previously shown by Kilian et al (2007) to be upregulated upon exposure to low temperature (4°C), suggesting a possibility that carbon assimilation could be an underlying mechanism inducing natural variations in photosynthesis at low temperature, and this might be mediated by trx. Interestingly, recent research showed that trx-h2 is essential for cold tolerance by upregulating cold-responsive genes in *Arabidopsis* (Park *et al*., 2021). We found trx-h1, but there is the possibility that different isoforms of trx affect chilling tolerance in different species.

The broadest range for the Chr 9 QTL spanned the region between 56.08-104.15cM (flanking markers, 2_01496 and 2_23951), and encompassed 1242 predicted coding regions (Table S9). This region contained several photosynthesis-related genes such as Mog1/PsbP/DUF1795-like photosystem II reaction center PsbP family protein (*Vigun09g156000 and Vigun09g204000*), subunit NDH-M of NAD(P)H: plastoquinone dehydrogenase complex (*Vigun09g160700*), ATP synthase epsilon chain *(Vigun09g163500*), photosystem II 22kDa protein (PsbS) (*Vigun09g165900*), ferredoxin-related (*Vigun09g220600*), photosystem II subunit X (*Vigun09g221400*), photosystem I light-harvesting complex gene 5 (*Vigun09g238500*), cytochrome *b6f* complex subunit (petM) (*Vigun09g241500*), plastocyanin (petE) (*Vigun09g257300*) and Photosystem II reaction center PsbP family protein (*Vigun09g263400*), as well as members of the thioredoxin superfamily (*Vigun09g154700, Vigun09g167600, Vigun09g224600, Vigun09g238200 and Vigun09g256300*) and thioredoxin M-type 4 (*Vigun09g156800*).

### The LIgHT: Illuminating Underlying Mechanisms, Not Just Candidate Genes

In this work, we explored stress-induced responses of a range of related, rapidly measurable photosynthetic processes in an RIL population of cowpea lines through the LIgHT approach. These responses reflect the genetically controlled variations in the control or regulation of photosynthesis. This approach is distinct from classical genetics, where mutations typically inactivate one or a few distinct enzymes in each genotype, leading to discrete loss of function phenotypes. Here, we may see combinations of effects that impact networks of processes are more likely to be adaptive.

Considering that the QTL regions in our study encompass hundreds of genes, we do not extensively explore the identities of specific, causative candidate polymorphisms. However, we have identified several interesting candidates, including structural and regulatory components of photosynthesis (Table. S6-9). In some cases, it is possible to identify the causative genetic components that underlie QTL or GWAS effects (e.g. (Caicedo *et al*., 2004; Roux *et al*., 2005)), but these cases are relatively few considering the number of published studies on genetic variation and quantitative trait locus (QTL) mapping, partly because of the low resolution of the genetic maps of most diversity panels (Roff, 2007; Miles and Wayne, 2008; Baxter 2020). Nevertheless, even at lower resolution, such genomic associations can be used to study mechanisms by applying LIgHT and guide plant breeding efforts and may provide important leads for specific genetic components that can be tested in future work.

More importantly, co-linkages (or lack thereof) from the LIgHT approach can be used to formulate and test scientific hypotheses, as we have demonstrated here, and thus give new insights into the processes in which evolution has modulated physiological responses. This approach makes comparisons across genotypes, emphasizing genetically controlled differences rather than the biophysical mechanisms per se. In other words, we observe how the genetic variations existing in a population “tweak” the mechanisms of photosynthesis. Key to this approach is the fact that each genotype in the population may have many combinations of smaller, quantitative effects that add up or interact to achieve altered responses. The statistical analyses of associations between the genetic components and measured parameters can give insights into the processes that control particular phenotypes. By comparing these associations across phenotypes, we can get further insights into how genetic variations affect the connections among related processes, i.e., which processes are potentially mechanistically or genetically linked to others.

Analysis of our cowpea RILs using time-resolved, high-throughput methods points to a model where important genetic control at the levels of the redox states of Q_A_ and *pmf*, which governs accelerated PSII damage, possibly due to the recombination reactions within PSII leading to singlet O_2_ production (Figure 7). We suggest that applying these methods to germplasm resources, such as mapping populations or diversity panels from diverse species, will reveal additional mechanisms of adaptation and will guide the breeding and engineering of photosynthesis for higher, more climate-resilient productivity.

## Abbreviations

LEF: linear electron flow
CEF: cyclic electron flow
PSI: photosystem I
PSII: photosystem II
Φ_II_: quantum efficiency of photosystem II
QTL: quantitative trait loci
NPQ: nonphotochemical quenching
LIgHT: Linkage Integration Hypothesis Testing
RILs: recombinant inbred lines
*pmf*: proton motive force
Chr: chromosome
NLM: nyctinastic leaf movements
Q_A_: primary quinone acceptor of PSII redox state of the PSII (Q_A_)
qL: a fraction of PSII centers in open states
ESTs: expressed sequence tags
MQM: Multiple QTL Mapping
KOG: EuKaryotic Orthologous Groups
KO: Kyoto Encyclopedia of Genes and Genomes
GO: Gene Ontology
DI: deionized water
HL: high light
CT: control temperature
LT: low-temperature
PCON: photosynthetic control
P_700_: primary electron donor of PSI
k_b6f_: rate constant for P_700_+ re-reduction k_b6f_
H^+^: proton
e^-^: electron
CB27: California Blackeye 27
IRAD: Institute de Recherche Agricole pour le Développement
DEPI: Dynamic Environmental Phenotype Imager
F_v_/F_m_: maximal PSII quantum efficiency
NPQt: non-photochemical quenching calculated using a theoretical (t) F_v_/F_m_ value
qIt: photoinhibition-related quenching calculated using a theoretical (t) F_v_/F_m_ value
qEt: energy-dependent quenching calculated using a theoretical (t) F_v_/F_m_ value
trx: thioredoxin
AGPase: ADP-glucose pyrophosphorylase

## Supplementary data

The following supplementary data are available at JXB online.

Table S1. List of Recombinant inbred line (RIL) parental crossings used for screening.

Table S2. Average temperatures experienced in cowpea fields.

Table S3. Experimental conditions for DEPI experiment.

Table S4. List of QTL intervals identified in photosynthetic parameters and NLM from DEPI.

Table S5. List of QTL intervals identified in photosynthetic parameters from MultispeQ.

Table S6. List of candidate genes in the minimum genetic range of QTL intervals on Chr 4.

Table S7. List of candidate genes in the maximum genetic range of QTL intervals on Chr 4.

Table S8. List of candidate genes in the minimum genetic range of QTL intervals on Chr 9.

Table S9. List of candidate genes in the maximum genetic range of QTL intervals on Chr 9.

Figure S1. Experimental design for growth and photosynthetic assays for QTL mapping and lincomycin treatment.

Figure S2. The photosynthetic response screening of RIL parental lines upon chilling stress.

Figure S3. Photosynthetic measurements of selected parental lines in the DEPI chamber over three days.

Figure S4. Hypothetical mechanistic contributors to photosynthetic responses at low temperature.

Figure S5. Histograms of photosynthetic parameters from DEPI across RIL lines taken at the middle of the third day (highest light intensity).

Figure S6. Significant changes and directionality of five photosynthetic parameters from DEPI in the low temperature (LT) compared to control conditions (CT).

Figure S7. Relative estimates of nyctinastic leaf movement (NLM).

Figure S8. Histograms of photosynthetic parameters from MultispeQ under both CT and LT conditions.

Figure S9. QTL analysis of photosynthetic parameters from DEPI in low-temperature conditions.

Figure S10. Time-resolved QTL analysis of five photosynthetic parameters under the control temperature (CT).

Figure S11. Time-resolved QTL analysis of five photosynthetic parameters under the low temperature (LT).

Figure S12. QTL analysis of photosynthetic parameters from MultispeQ in the control condition.

Figure S13. QTL analysis of photosynthetic parameters from MultispeQ in low temperature condition.

Figure S14. Time-resolved QTL analysis of NLM (or relative tip-to-tip distance) under both CT and LT conditions from DEPI chamber experiments.

Figure S15. The Daisy plots show associations for selected QTL intervals of photosynthetic and non-photosynthetic parameters from DEPI and MultispeQ in CT at Chr 4, 59.04-64.45 cM (A) and Chr 9, 86.93-104.15 cM (B) Chr 8, 34.73-35.34 cM (C) Chr 10, 20-30.7cM (D).

Figure S16. Time course for the appearance and disappearance of the QTLs of five photosynthetic parameters at the selected two loci, Chrs 4 and 9.

Figure S17. Effect plots of identified QTLs in Chrs 4 and 9 for Φ_II_ and qIt at 1.5 hr prior to the end of Day 3 (206 µmol, m^-2^, s^-1^).

Figure S18. Box plots for three allele groups of identified QTLs in Chrs 4 and 9 for Φ_II_ and qIt at 1.5 hr prior to the end of Day 3 (206 µmol, m^-2^, s^-1^).

Figure S19. Box plots for four allele groups of identified QTLs in Chrs 4 and 9 for Φ_II_ and qIt at 1.5 hr prior to the end of Day 3 (206 µmol, m^-2^, s^-1^).

Figure S20. Relationships between photosynthetic responses grouped by different combinations of alleles for the identified QTLs in Chrs 4 and 9 under both CT and LT conditions (CT: open circles, LT: closed circles).

Figure S21. Significance matrices (p-values in each box) of five photosynthetic parameters for four allele groups are shown in Fig 6 under CT (A-E) and LT (F-J) conditions.

## Acknowledgments

The authors thank Professors Douglas Schemske (Michigan State University), Christopher Oakley (Purdue University), Tim Close (UC Riverside) and Dr. Tom P.J.M. Theeuwen (Jan IngenHousz Institute) for helpful discussions on QTL analyses and Professors Christoph Benning (Michigan State University), Michael Thomashow (Michigan State University) and Patrick Horn (University of North Texas) for discussions on chilling effects. The authors thank Mr. David Hall and Mr. Oliver L. Tessmer for their technical assistance. Above all else, we would like to express our deepest gratitude to Dr. Wellington Muchero for his invaluable insights into this research. In memory of Dr. Wellington Muchero, we dedicate this work to the memory of our colleague and friend. We will miss his wisdom, his humor and his support.

## Author contributions

DH and DMK conceived the idea and interpreted the data. DH conducted the experiments and analyzed data. DH and DMK wrote the manuscript. BLH and PAR provided cowpea genetic materials, marker data, and agronomy guidance. All authors helped revise the manuscript. All authors approved the final manuscript.

## Data availability

The data supporting the research is available in the paper and supplementary materials, which are published online. Python codes are available in the GitHub repository at https://github.com/DongheeHoh/LIgHT.

